# Phosphatase SHP2 pathogenic mutations enhance activity by altering conformational sampling

**DOI:** 10.64898/2025.12.12.694068

**Authors:** Andrew W. Glaser, Ricardo A. P. de Pádua, Adedolapo M. Ojoawo, Camille Sullivan, Dorothee Kern

**Author notes:** These authors contributed equally (Andrew W. Glaser, Ricardo A. P. de Pádua, Adedolapo M. Ojoawo).

## Abstract

SH2 domains are critical mediators of cellular signaling, although the molecular mechanisms by which they bind their phosphopeptide ligands remain incompletely understood. We investigate the atomic mechanisms underlying both healthy regulation and dysregulation of the human protein tyrosine phosphatase SHP2, a key regulator of cellular signaling. While most pathogenic mutations cluster near the PTP/N-SH2 interface, the E139D and T42A mutations are located within the regulatory SH2 domains, and their mechanisms of dysregulation remain controversial. The T42A mutation in the N-SH2 domain paradoxically increases phosphotyrosine-peptide binding affinity despite disrupting the hydrogen bond of T42 to the phosphoryl group, a puzzling contradiction that remains unresolved. We find that the T42A mutation shifts the conformational ensemble of peptide-bound N-SH2 toward a zipped β-sheet state and suppresses millisecond conformational exchange, supporting a model in which enhanced stabilization of the zipped conformation contributes to hyperactivation. This conformational shift provides a structural rationale for the increased affinity of T42A and helps reconcile previously conflicting models of peptide-induced SHP2 activation. By integrating X-ray ensemble refinement with NMR relaxation, our work illustrates how complementary structural and dynamic approaches can uncover regulatory mechanisms in SHP2 and may inform broader principles of SH2-mediated phosphopeptide recognition.

**Significance Statement:** Here, we characterize how two pathogenic SH2-domain mutations alter SHP2 regulation and lead to hyperactivation. We identify a previously unobserved apo conformation of the N-SH2 domain in which Tyr66 occludes the peptide-binding cleft, indicating that a conformational change is required for full binding of activating phosphopeptides. Our data suggest that the T42A mutation shifts the equilibrium toward a zipped central β-sheet state in the peptide-bound N-SH2 domain as the most likely model underlying the measured 10-fold increased binding affinity. These results help clarify the structural basis for SHP2 regulation and illustrate how conformational dynamics shape SH2-phosphopeptide recognition.

## Introduction

SHP2, a protein tyrosine phosphatase, is central to numerous signaling pathways. Mutations in *PTPN11*, which encodes SHP2, are linked to a spectrum of diseases, including Noonan syndrome, LEOPARD syndrome, and cancers like juvenile myelomonocytic leukemia.(1–3) SHP2 comprises a catalytic protein tyrosine phosphatase (PTP) domain, two regulatory SH2 (Src homology 2) domains (N-SH2 and C-SH2), and an unstructured C-terminal tail.(4, 5) Unperturbed, SHP2 exists in equilibrium between an active, open (10%), and an inactive, closed conformation (90%) in which the N-SH2 occludes the PTP active site (**Fig. 1A**).(6–8)

**Fig. 1.**
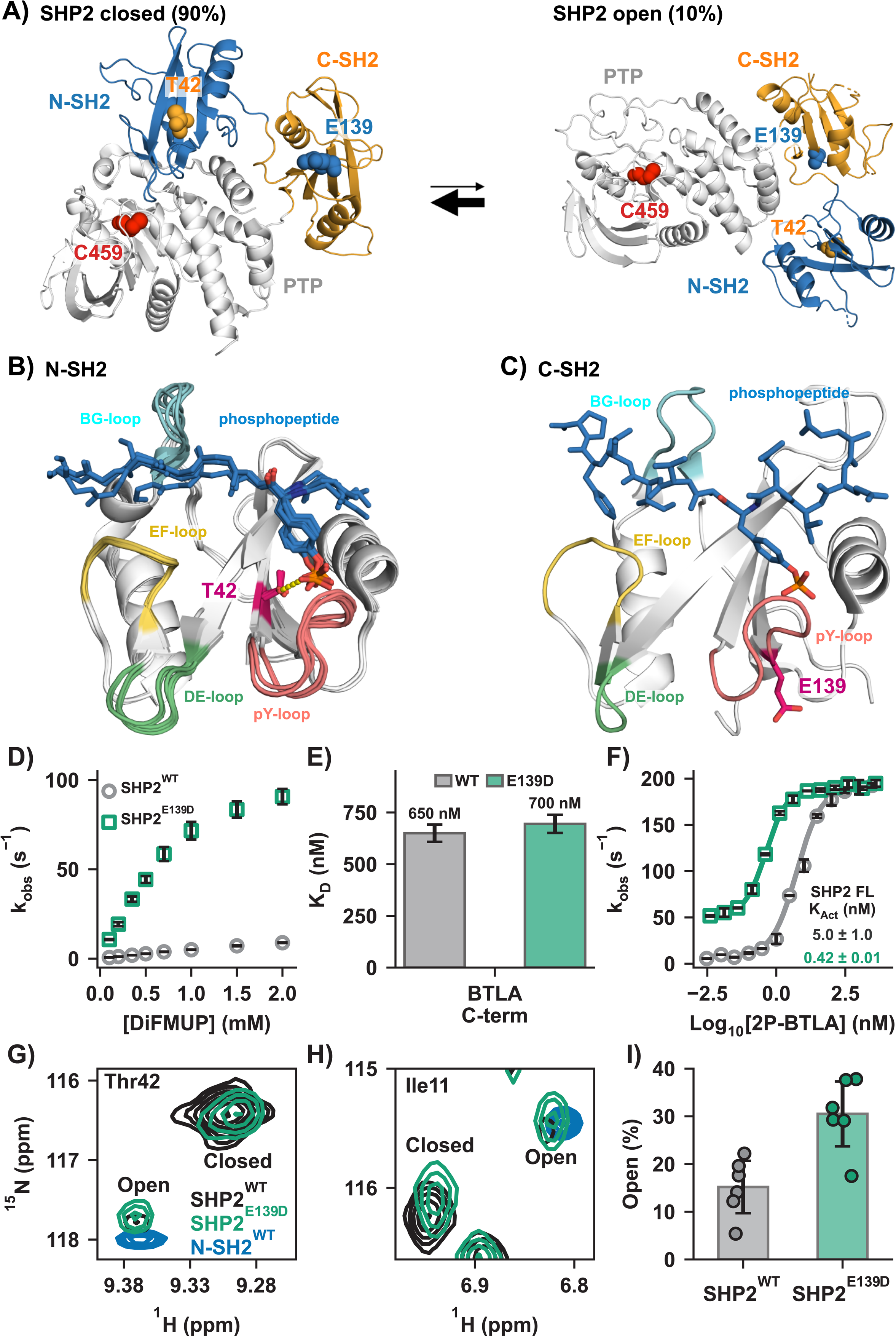
Structure and activation of SHP2 WT and E139D mutant. **(A)** SHP2 exists in equilibrium between the auto-inhibited, closed form (left) (PDB code: 4DGP) and active, open form (right). Activation shifts this equilibrium to the open form, in which the N-SH2 domain disassociates from the PTP domain, rendering the active site accessible to the substrate (PDB code: 6CRF). **(B)** Interaction of Thr42 (magenta) in the N-SH2 domain with various phosphopeptide ligands (blue), as visualized in several crystal structures (PDB codes: 6ROY, 6ROZ, 4QSY, 5DF6, 3TZ0). The H-bond between the Thr42 hydroxyl and the phosphate group is shown as yellow dotted line. **(C)** C-SH2 domain of wildtype SHP2 bound to a phosphopeptide ligand (blue); Glu139 side chain (magenta) is oriented away from the C-SH2 p-Tyr binding site (PDB code: 6R5G). **(D)** Representative K_M_ curves highlight the increase in basal activity for SHP2^E139D^ (green) compared to SHP2^WT^ (gray). **(E)** The phosphopeptide binding affinity remains unchanged with the E139D mutation as shown by unaltered dissociation constant (K_D_) of C-SH2 to 1P-BTLA_C-term_ using isothermal titration calorimetry (ITC). **(F)** Representative K_Act_ curves of SHP2^WT^ (grey) and SHP2^E139D^ (green) using 2P-BTLA. **(G, H)** Direct detection of the major (closed) and minor (open) states of SHP2 by NMR. ^1^H-^15^N TROSY HSQC spectra of perdeuterated SHP2^WT^ (black) and SHP2^E139D^ (green). ^1^H-^15^N TROSY HSQC spectra of SHP2 N-SH2 (blue) is shown as a marker for the open conformation. **(I)** Quantitation of major (closed) and minor (open) peak volume for SHP2^WT^ and SHP2^E139D^. Crucially, the E139D mutation shifts the open/closed equilibrium of SHP2 from 15 ± 6% for SHP2^WT^ to 31 ± 7% for SHP2^E139D^.

SH2 domains recognize and bind phosphotyrosine-containing sequences, mediating protein-protein interactions essential for signal transduction.(9–11) In SHP2, PTP activity is allosterically regulated by bis-phosphotyrosine motifs binding to each SH2 domain, stabilizing the protein’s open form.(12, 13) SHP2 activators include B- and T-lymphocyte attenuator (BTLA), which recruits SHP2 to suppress immune response, and GRB2-associated-binding protein 1 (GAB1), a soluble docking protein that activates SHP2 in various signaling pathways.(14–19) Different activators enable SHP2 to function selectively across multiple signaling pathways.(20)

Many disease-related SHP2 mutations disrupt the N-SH2/PTP interface, promoting the active conformation.(1, 21, 22) In contrast, regulatory SH2 domain mutations, less studied, offer insight into SH2/phosphopeptide binding and activation. Our study examines two such pathogenic mutations, T42A and E139D **(Fig. 1B, C)**. While Thr42 and Glu139 are poorly conserved among human SH2 sequences (**Fig. S1**), they are highly conserved among SHP2 sequences across species (**Fig. S1**). Both mutations upregulate PTP activity.(23–25)

E139D increases SHP2’s basal activity and affinity to activating peptides, but its molecular mechanism remains unclear.(26–29) Located in the C-SH2 p-Tyr binding pocket, E139D is distant from the N-SH2/PTP interface, unlike the stereotypical mutations that alter the equilibrium to favor the open state (**Fig. 1C**).

The T42A mutation enhances N-SH2 binding affinity for phosphopeptides, although its molecular mechanism is unknown.(26, 27, 30) Crystallography reveals that the hydroxyl of Thr42 forms a hydrogen bond with the phosphoryl group of the peptides’ p-Tyr (**Fig. 1B**).(31) Removing this interaction by the T42A mutation could naively be interpreted as resulting in weakened binding, but the energetic contribution of this interaction has not been measured. However, studies report that T42A increases affinity for phosphorylated activators, ranging from a few-fold to 50-fold, depending on the phosphopeptide sequence.(29, 31) Molecular dynamics (MD) simulations have proposed various mechanisms for peptide binding, but failed to explain this increased affinity for T42A.(28, 32–35)

To resolve this, we dissect the mechanisms of these perplexing mutations through biochemical and biophysical experiments. We found that the E139D mutation shifts the open/closed equilibrium to 35% open without altering the affinity of p-Tyr peptides.(29) We solved a high-resolution X-ray structure of the apo N-SH2 domain that presented a novel conformation; combining this with ensemble refinement of X-ray structures of the peptide-bound forms for both WT and T42A and NMR dynamics, we discovered a novel multi-step binding mechanism of p-Tyr-containing peptides that explains the increased peptide affinity for T42A. These results highlight the power of combining crystallography with NMR dynamics experiments. Our activation model likely applies broadly to SH2 domain peptide binding.

## Results

### The E139D mutation upregulates PTP activity by increasing SHP2’s open population

To confirm the previously reported effects of the E139D mutation,(23, 26, 27) we evaluated its impact on basal activity and phosphopeptide binding. SHP2^E139D^ exhibited significantly higher basal activity relative to SHP2^WT^, measured via hydrolysis of the artificial substrate DiFMUP, in which dephosphorylation was tracked through absorbance of the dephosphorylated product DiFMU (**Fig. 1D**). Isothermal titration calorimetry (ITC) showed that C-SH2^WT^ and C-SH2^E139D^ bind the singly phosphorylated BTLA_C-term_ motif (1P-BTLA_C-term_) (**Fig. 1E** and **Fig. S2**, **S3**) with similar affinities, consistent with previous studies.(29) SHP2^E139D^ activation by the bis-phosphotyrosine-containing activator BTLA (2P-BTLA) displayed a more-than-10-fold tighter K_Act_ than SHP2^WT^ (**Fig. 1F**).

These findings suggest that E139D increases basal enzyme activity (and the apparent K_Act_) by shifting the equilibrium towards the open state. To directly monitor this shift, we collected ^1^H-^15^N TROSY-HSQC spectra of SHP2^WT^ and SHP2^E139D^. Peak duplication for several amides indicated slow interconversion between distinct conformations, consistent with previously reported SHP2 opening and closing rates.(7) Using the isolated N-SH2 domain as an open-state marker, we identified six residues as reporters of the open/closed equilibrium (**Fig. 1G-H**). Peak volume quantitation estimated the population of open SHP2^WT^ to 15 ± 6%, which is within experimental uncertainty of our previously reported result of 10.4 ± 1.1%, despite methodological differences.(7) We measured that the E139D variant is roughly 2-fold more open than the WT protein (**Fig. 1I**) with an open population of 31 ± 7%. The 2-fold increase in open population, measured in the absence of DiFMUP, is amplified when coupled with substrate binding at the saturating concentrations used in the activation assays, resulting in a 10-fold increase in activity and therefore a 10-fold tighter K_Act_ for SHP2^E139D^ **(Fig. 1F)**. We note that this same phenomenon was observed for the SHP2^E76D^ mutant which also caused a 2-fold increase in the open population.(7)

In an effort to structurally explain the increase in the open population by the E139D mutation we superimposed crystal structures of SHP2^WT^ and SHP2^E139D^ to detect differences in the local structure (**Fig. S4**). We only used crystal structures of the closed state without ligands, including several for SHP2^WT^ and one for SHP2^E139D^.(27) We noticed subtle structural differences in the region near residue 139 due to the mutation. In SHP2^WT^, we observe a range of conformational substates in the N-SH2/C-SH2 linker, particularly for His116. In contrast, His116 adopts a unique conformation in SHP2^E139D^; the side chain of His116 is oriented away from the PTP/C-SH2 interface, potentially destabilizing the closed state. Given the lack of electron density observed around Glu139 in the open conformation of SHP2^E76K^, it is unlikely that Asp139 stabilizes the open conformation seen in the SHP2^E76K^ open crystal structure.(36) Alternatively, the AlphaFold 2 predicted open conformations of 2P-BTLA bound SHP2^WT^ and SHP2^E139D^ suggest an interaction between residue 139 and Arg5 that may stabilize the open state (**Fig. S4**). Because the E139D mutation increases the open population by only 2-fold, the accompanying ΔΔG of-0.41 kcal/mol is less than the free energy of a hydrogen bond, making it hard to decipher the correct structural mechanism.

### The T42A mutation increases phosphopeptide affinity without altering the open/closed equilibrium

SHP2^T42A^ and SHP2^T42S^ exhibited basal activity comparable to SHP2^WT^, confirming no equilibrium shift (**Fig. 2A**). However, N-SH2^T42A^ showed a 10-fold increase in affinity for 1P-BTLA_N-term_ (**Fig. 2B, S5**) and an 8-fold increase for 1P-GAB1_N-term_, while N-SH2^T42S^ displayed intermediate affinities. Enzyme assays on the full-length proteins with 2P-BTLA showed a 10-fold smaller K_Act_ for SHP2^T42A^ compared to SHP2^WT^ (**Fig. 2C**), directly due to the 10-fold higher affinity of the N-SH2^T42A^ domain.

**Fig. 2.**
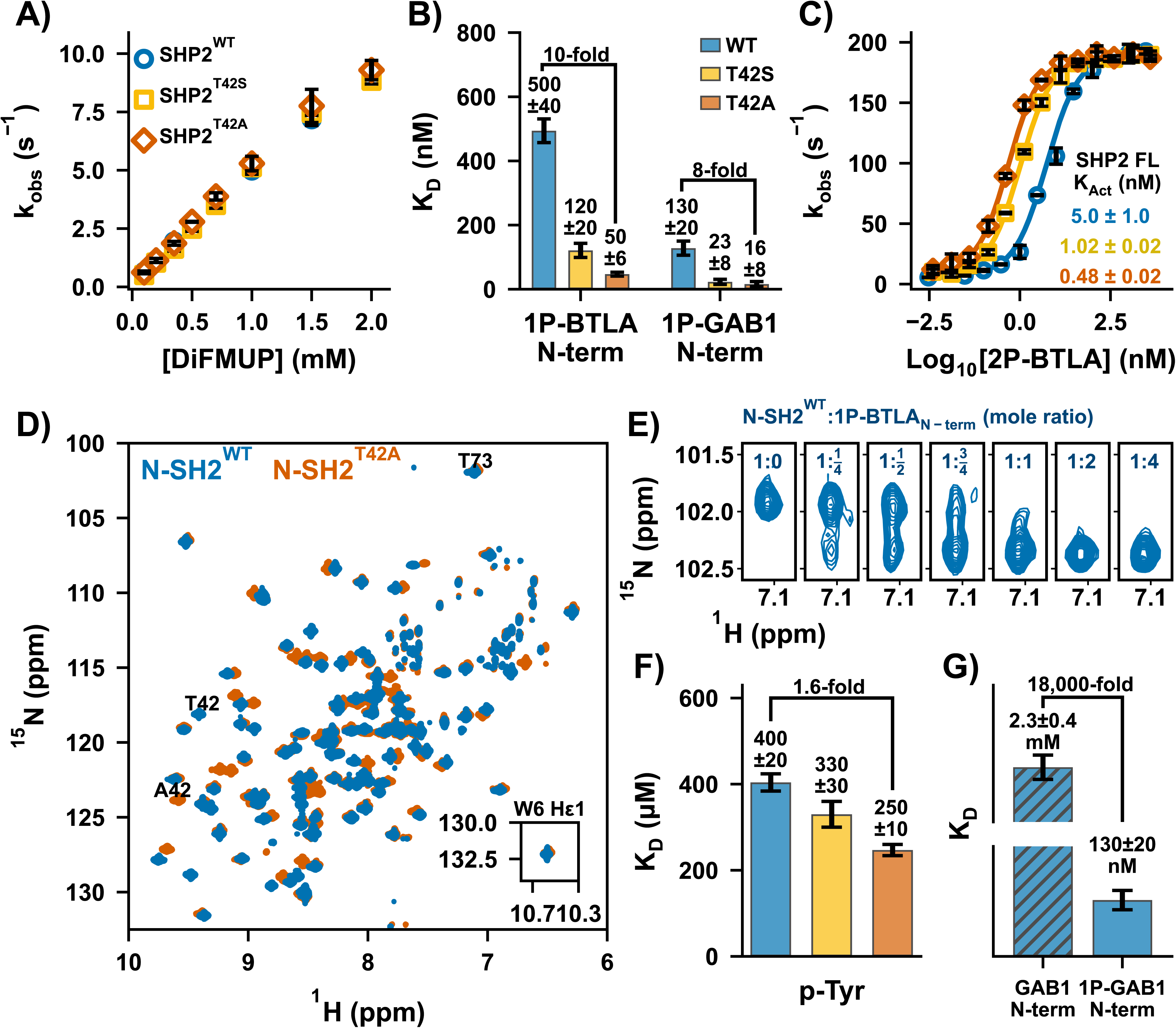
Molecular mechanism of T42 mutations in N-SH2 domain for SHP2 upregulation. (A) Representative K_M_ curves reveal identical basal activities for SHP2^T42S^ and SHP2^T42A^ compared to SHP2^WT^. **(B)** N-SH2^T42A^ variant showcases a 10- and 8-fold increase in binding affinities to two phosphopeptide ligands: 1P-BTLA_N-term_ and 1P-GAB1_N-term_, respectively. **(C)** Representative K_Act_ curves of SHP2^WT^, SHP2^T42S^, and SHP2^T42A^ using 2P-BTLA. **(D)** ^1^H-^15^N TROSY HSQC spectra of N-SH2^WT^ and N-SH2^T42A^. **(E)** Titration of unlabeled 1P-BTLA_N-term_ phosphopeptide to ^15^N labeled N-SH2^WT^, where peak duplication upon binding was observed. **(F)** p-Tyr binding affinity to N-SH2 variants. K_D_ values extracted from chemical shift perturbations for N-SH2^WT^, N-SH2^T42S^, and N-SH2^T42A^ upon titration with p-Tyr. **(G)** Unphosphorylated GAB1_N-term_ binds 18,000-fold weaker to N-SH2^WT^ compared to the phosphorylated peptide.

### Dissecting the binding contribution of p-Tyr

To investigate the mechanism underlying N-SH2^T42A^’s enhanced binding affinity, we analyzed ^1^H-^15^N HSQC spectra of the apo and bound N-SH2 domains. Significant chemical shift perturbations (CSPs) were observed near the mutation site, with minor CSPs throughout the domain, indicating a global effect on protein conformation (**Fig. 2D, S6**). Titration of 1P-BTLA_N-term_ phosphopeptide into N-SH2^WT^ or N-SH2^T42A^ showed peak duplication between the apo and bound states, indicating a slow exchange binding process consistent with the measured nM K_D_. Over-titration of peptide revealed no further peak shifts (**Fig. 2E, S7**). However, rapid processes impacting peptide binding might be overlooked in this simple NMR experiment.

Given the paradoxical result of increased affinity of N-SH2^T42A^ by removing the H-bonding T42 hydroxyl to p-Tyr, we directly measured p-Tyr affinity to all three variants by an NMR titration.(34) Binding occurred in the fast-exchange regime, leading to peak shifts. Fitting CSPs as a function of p-Tyr concentration yielded K_D_ values of 400 µM, 330 µM, and 250 µM for N-SH2^WT^, N-SH2^T42S^, and N-SH2^T42A^, respectively. While N-SH2^T42A^ showed a modest 1.6-fold increase in p-Tyr affinity, this failed to explain the 10-fold increase in overall phosphopeptide binding affinity (**Fig. 2F, S8**).

N-SH2^WT^ binds p-Tyr with a K_D_ of ∼400 µM, much weaker than the phosphopeptides 1P-BTLA_N-term_ and 1P-GAB1_N-term_, (500 nM and 130 nM, respectively), demonstrating a 1000-5000-fold affinity increase due to additional peptide interactions. Strikingly, the unphosphorylated GAB1_N-term_ peptide also bound extremely weakly, with a K_D_ of 2.3 mM – approximately 18,000 times weaker than its phosphorylated counterpart (**Fig. 2G, S8**). This stark contrast suggests that T42A’s effects extend beyond merely enhancing p-Tyr binding; they influence the binding of the entire phosphopeptide in a manner that cannot simply be explained by the sum of its parts.

### Crystal structures reveal unique binding conformations that stabilize the peptide-bound ensemble for N-SH2^T42A^

To investigate the cooperative mechanism underlying the enhanced binding, we solved crystal structures of each SH2 domain in their apo forms and bound to 1P-GAB1_N-term_ (**Table S1**). We uncovered a previously unreported feature of the apo N-SH2 that provides novel insight into the binding mechanism: In our apo structure, Tyr66 blocks the binding cleft due to steric clashes with C-terminal residues adjacent to p-Tyr, while the phosphotyrosine binding loop is already compatible with p-Tyr binding (**Fig. 3A**). This suggests a multi-step binding process in which p-Tyr initially binds, followed by a conformational shift, repositioning Tyr66 to accommodate the full peptide, a feature present in both variants. Since static structures do not fully capture conformational flexibility, we refined ensemble models to better represent solution dynamics. The ensemble models revealed greater diversity for N-SH2^WT^ than N-SH2^T42A^ most prominent in the peptide-bound states (**Fig. 3B-E**).(37)

**Fig. 3.**
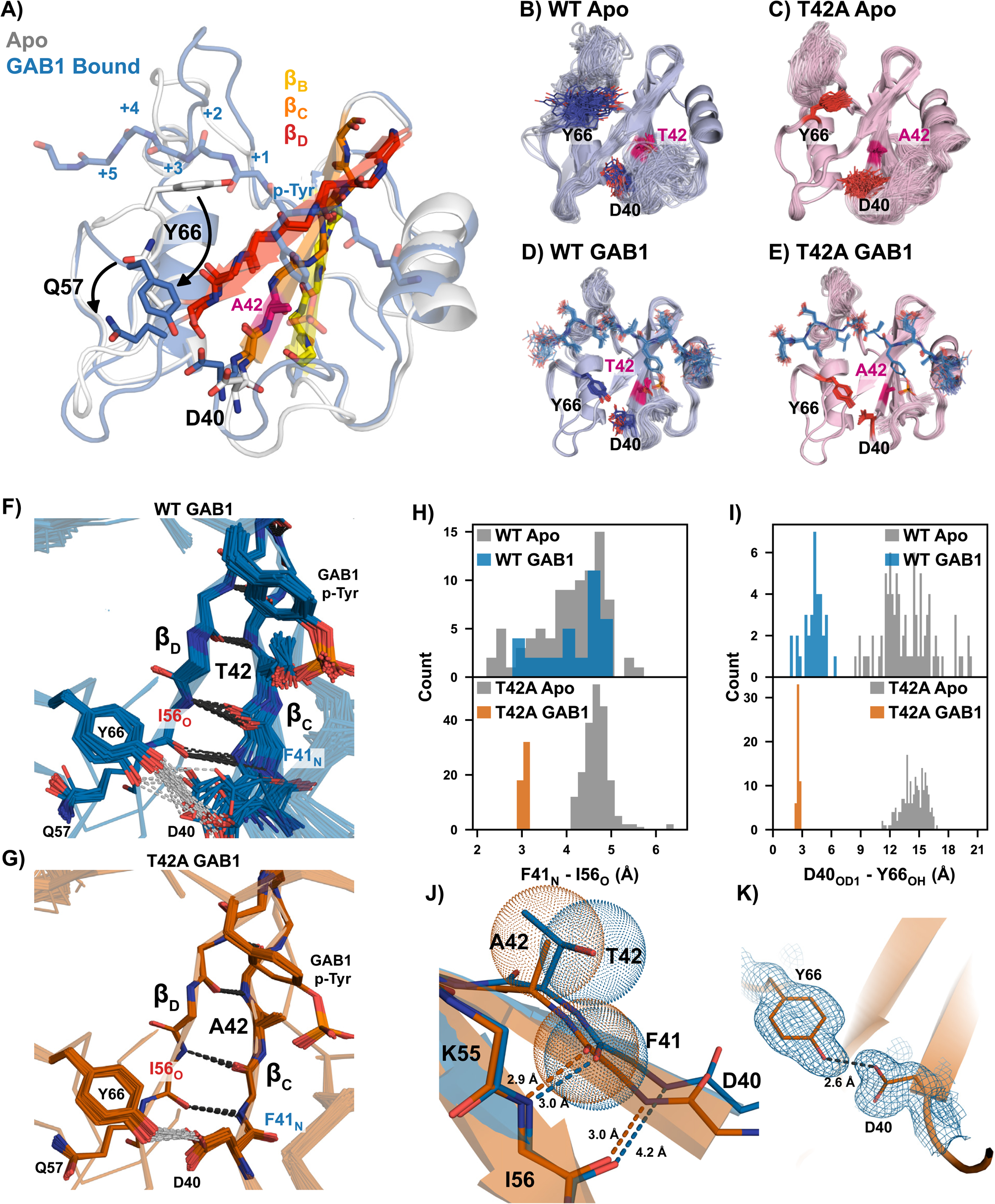
Structural rationale of altered T42A phosphopeptide binding affinity. **(A)** Apo (white) and 1P-GAB1_N-term_-bound (labeled GAB1 in figure) (blue) structures of the N-SH2^T42A^. The apo form of the protein is incompatible with complete peptide binding due to Tyr66 blocking the hydrophobic binding cleft, which flips out upon peptide binding (arrow). The β-strands essential for correlated conformational changes for peptide binding are labeled. Ensemble refinement crystal structures of apo N-SH2^WT^ **(B)** and N-SH2^T42A^ **(C)**; and 1P-GAB1_N-term_-bound forms **(D)** and **(E)**. **(F,G)** Zoom in of the beta strands β_C_ and β_D_ of 1P-GAB1_N-term_-bound N-SH2^WT^ **(F)** and N-SH2^T42A^ **(G)**. Beta strand unzipping is observed for N-SH2^WT^ **(F)**, while N-SH2^T42A^ fully populates zipped β-strands being further stabilized by hydrogen bonding between the side chains of Y66 and D44 **(G)**. **(H)** Quantification of Ile56_O_ - Phe41_N_ and **(I)** Tyr66 - Asp40 distance in the apo and 1P-GAB1_N-term_-bound forms of N-SH2^WT^ and N-SH2^T42A^ in each ensemble model. **(J)** Strand-strand interaction formed by the interaction of Phe41_N_ and Ile56_O_. The bulkier threonine sidechain prevents optimal strand-strand interaction as shown by van der Wals radii. **(K)** Formation of a short, strong hydrogen bond between Tyr66 and Asp40 in the N-SH2^T42A^ 1P-GAB1_N-term_-bound structure.

Peptide binding to N-SH2 requires repositioning of Tyr66 to accommodate the full ligand. Binding of 1P-GAB1_N-term_ stabilizes both variants, but N-SH2^T42A^ exhibits less flexibility than N-SH2^WT^, particularly in the p-Tyr loop **(Fig. 3D,E).** Importantly, 1P-GAB1_N-term_-bound N-SH2^WT^ adopts two conformations. In most structures, the ends of beta strands β_C_ and β_D_ are separated, indicated by a 4 – 5 Å distance between Phe41_N_ and Ile56_O_, precluding hydrogen bonding, which we call ‘unzipped’ conformation. A minority of 1P-GAB1-bound N-SH2^WT^ structures shows a narrower Phe41_N_ - Ile56_O_ distance (2.7 – 3.3 Å), allowing for hydrogen bonding and extending beta strand β_D_ (**Fig. 3F, H**, ‘zipped’ conformation).

Strikingly, the bound form of N-SH2^T42A^ exclusively occupied the zipped conformation (**Fig. 3G,H**). The bulkier side chain of Thr42 in N-SH2^WT^ causes steric clashes that disrupts optimal β_C_/β_D_ interaction, favoring the spread conformation. In contrast, the smaller Ala42 side chain in N-SH2^T42A^ alleviates this clash, stabilizing an extended, zipped beta strand (**Fig. 3J**). Moreover, the zipped conformation enables an additional stabilizing hydrogen bond between the Tyr66 hydroxyl and the Asp40 carboxylate (**Fig. 3F,G,I,K**). Notably, this hydrogen bond has an average bond length of 2.6 Å in N-SH2^T42A^, which is shorter than the typical range of hydrogen bond lengths.(38) Short hydrogen bonds (SHBs) are common between Aspartate and Tyrosine, providing additional stability to the N-SH2^T42A^ peptide-bound complex.

Interestingly, in both apo states, Gln57 in β_D_ sterically blocks the Tyr66-Asp40 interaction; requiring a conformational shift before the SHB can form. Furthermore, the unzipped β strand conformation is dominating (**Fig. 3H**).

Analysis of the bound N-SH2^T42A^ structural ensemble revealed also enhanced p-Tyr loop/phosphotyrosine interactions over N-SH2^WT^. The sidechains of Ser34 and Ser36, along with the backbone of Lys35, are positioned closer to the phosphate oxygens in N-SH2^T42A^ (**Fig. S9**). These improved interactions agree with the tighter binding of p-Tyr to N-SH2^T42A^ measured by NMR **(Fig. 2F)** and with increased enthalpic interactions observed in ITC experiments for 1P-GAB1_N-term_ and 1P-BTLA_N-term_ peptides (**Fig. S5**). The more negative enthalpy change (ΔH) in T42A indicates stronger or more numerous stabilizing interactions in the mutant complex.

Finally, the crystal structure of N-SH2^WT^ bound to p-Tyr alone revealed a striking result (**Fig. S10**): only the unzipped conformation was observed. However, as the entire peptide binds, the zipped conformation becomes more populated, with a full shift to the zipped conformation in N-SH2^T42A^. In the p-Tyr-bound structure, Tyr66 still blocks the peptide binding cleft and Gln57 prevents the formation of a stabilizing Tyr66-Asp40 hydrogen bond. The crystallographic p-Tyr-bound unzipped conformation demonstrates that phosphotyrosine can engage the binding pocket prior to β-sheet closure. This observation indicates that p-Tyr recognition can precede full peptide docking, without implying a specific binding pathway.

### NMR relaxation reveals timescales of conformational sampling causing T42A hyperactivation

We used ^15^N R_1_, R_2_, and heteronuclear NOE (hetNOE) relaxation techniques to assess whether the ‘zipped’ and ‘unzipped’ conformations found in X-ray structures exchange on the picosecond to nanosecond (ps-ns) timescale. If so, changes in hetNOE values would be detectable. These experiments that probe ps-ns timescale motions were particularly relevant to perform as all previous models for activator binding to the N-SH2 domain proposed mechanisms grounded in this timescale.(32, 33)

Our hetNOE results indicate a generally rigid backbone in both the apo and 1P-GAB1_N-term_-bound forms of the N-SH2 variants (**Fig. 4A, Fig. S11**), with increased flexibility in the p-Tyr, DE, and EF loops in the apo form that rigidify upon peptide binding. This aligns with our findings from crystallography and previous studies(34), where phosphopeptide binding stabilizes these loops. Notably, hetNOE values for 1P-GAB1_N-term_-bound N-SH2^WT^ and N-SH2^T42A^ were similar, indicating that the ‘zipped’ to ‘unzipped’ conformational exchange does not occur on the ps-ns timescale.

**Fig. 4.**
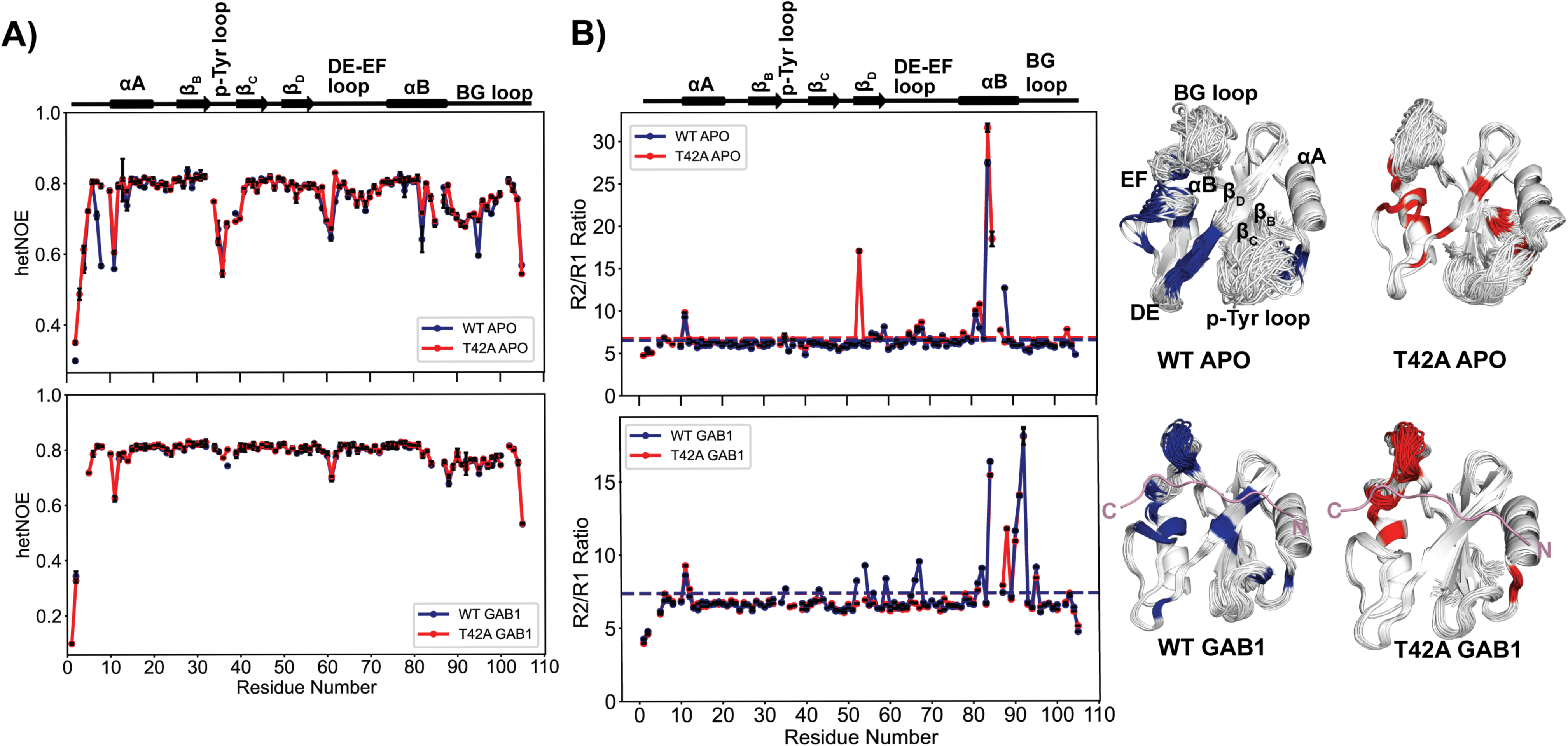
^15^N backbone NMR dynamics to probe picosecond to nanosecond (ps-ns), and microsecond-to-millisecond (µs-ms) motions. **(A)** Overlay of ^15^N- heteronuclear NOE (hetNOE) data comparing the apo and the 1P-GAB1_N-term_-bound (denoted GAB1 in figure labels) forms of N-SH2^WT^ and N-SH2^T42A^. Both proteins show very similar values, indicating that the ps-ns motions are very similar. **(B)** ^15^N R_2_/R_1_ ratios for the apo and 1P-GAB1_N-term_-bound forms of N-SH2^WT^ and N-SH2^T42A^. Horizontal lines denote a cutoff of 1 standard deviation above the mean R_2_/R_1_ ratio, indicative of µs-ms motions. Residues above the threshold are color-coded in the ensemble structures on the right and hint at a difference on this timescale between mutant and WT for the 1P-GAB1_N-term_-bound forms.

R_2_/R_1_ ratio plots (**Fig. 4B, Fig. S12**) reveal elevated R_2_ values in the β_D_ strand of both apo forms, and the 1P-GAB1_N-term_-bound N-SH2^WT^, while for 1P-GAB1_N-term_-bound N-SH2^T42A^, elevated R_2_ values are restricted only to the BG loop. Elevated R_2_ values for N-SH2^WT^ are consistent with a conformational exchange between ‘zipped’ and ‘unzipped’ conformation on the µs-ms timescale. The absence of elevated R_2_ values for bound N-SH2^T42A^ indicates that no such exchange occurs, suggesting that the beta sheet is locked into the zipped conformation. Common elevated R_2_ values in the BG loop for both bound N-SH2 variants suggest a separate µs-ms conformational process coupled to 1P-GAB1_N-term_ binding.

To directly measure the zipping/unzipping exchange timescales, we employed ^15^N-CPMG relaxation dispersion(39) and ^15^N-CEST(40), sensitive to µs-ms conformational processes. We extracted exchange rates and populations by fitting residue-specific relaxation data to a two-site model. Analysis of the 1P-GAB1_N-term_-bound N-SH2^WT^ CEST data revealed two groups based on their exchange rates (*k_ex_*) (**Fig. 5A-B, Fig. S13**); 28 residues with an exchange rate (*k_ex_ = k_AB_ + k_BA_*) = 104 ± 8.5 s^-1^, *pB* = 11 ± 0.3 %, while two residues (Leu43 and Ile54) showed a slightly slower exchange rate of 45 ± 5 s^-1^, *pB* = 16 ± 0.6 % (**Table S2**). Interestingly, residues with CEST profiles were in the β_C_-β_D_ strands but extended beyond these regions (**Fig. 5B**).

**Fig. 5.**
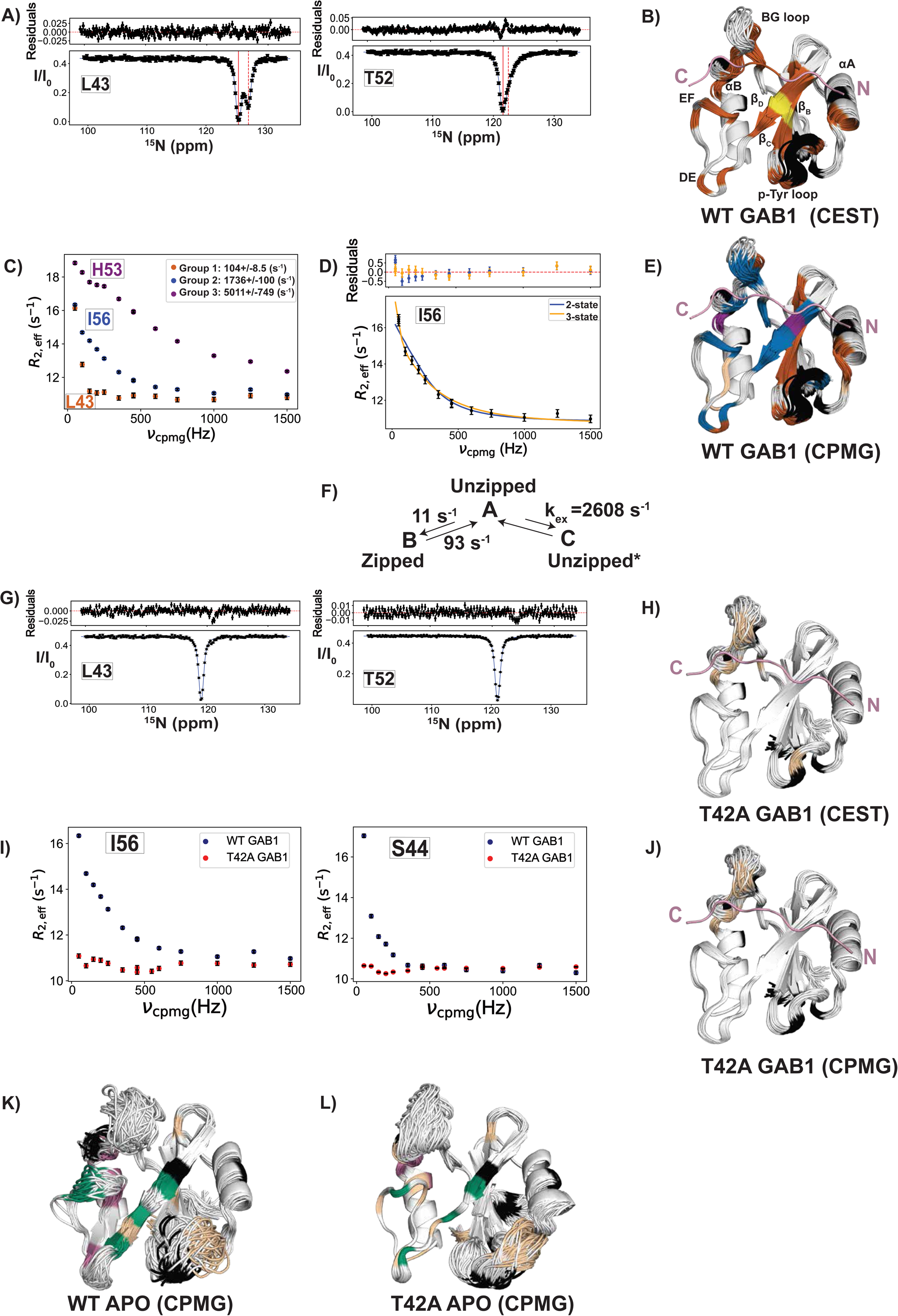
^15^N backbone NMR dynamics reveal ms timescale motion in peptide-bound forms likely responsible for increased peptide affinity of N-SH2^T42A^. **(A)** Representative ^15^N-CEST profiles of residues located in β_C_ (Leu43) and β_D_ (Thr52) for 1P- GAB1_N-term_-bound (denoted GAB1 in figure labels) N-SH2^WT^ fitted to a two-state exchange. The solid black line is the fit, the solid and dashed red lines indicate the major and minor state positions, respectively. **(B)** Data in **A** were globally fitted into two groups: *k_ex_ (k_AB_+k_BA_*) =104 ± 8.5 s^-1^, *pB* = 11 ± 0.3 % (brown), and *k_ex_ =* 45 ± 5 s^-1^, *pB* = 16 ± 0.6 %, (yellow), plotted onto our ensemble X-ray ensemble structures. **(C)** Representative ^15^N CPMG relaxation dispersion curves for 1P-GAB1_N-term_-bound N-SH2^WT^ from the three distinct groups of residues, each fitted to a two-state exchange model. The color of each group in the dispersion curves matches the color in **E. (D)** Six of group 2 residues in **C** needed to be fitted to a three-state model (scheme **F**), and a representative overlay of 2-state and 3-state fittings is shown. To obtain the fit, *k_ex,AB,_ a*nd Δω_AB_ were fixed using CEST parameters (*k_ex,AB_ =* 104 ± 8.5 s^-1^), yielding a *k_ex_,_AC_ =* 2600 ± 100 s^-1^, and underdetermined *pB*, *pC*. Ile54 which has a *k_ex,AB_ =* 45 ± 5 s^-1^ from CEST data, was separately fitted to a 3-state model, yielding a *k_ex_,_AC_*= 1830 ± 400 s^-1^, and underdetermined *pB, pC* (see **Fig. S15** for all profiles). Two-state fit (blue) is shown for comparison and fails to account for low-frequency deviations caused by the slow process detected by CEST. **(E)** X-ray ensemble structure of 1P-GAB1_N-term_-bound N-SH2^WT^ color-coded by the three groups from CPMG two-state fitting: group 1: *k_ex_* = 104 ± 8.5 s^-1^, *pB* = 11 ± 0.3 % (brown), group 2: *k_ex_ =* 1740 ± 100 s^-1^, underdetermined *pB* (blue), group 3: *k_ex_ =* 5000 ± 750 s^-1^, underdetermined *pB* (magenta). Tan-colored residues don’t fit into a group. **(F)** Final model explaining relaxation data depicting the conformational dynamics between the unzipped (A, major) and zipped (B, minor) states, as well as a more flexible unzipped minor state (C, denoted by unzipped*) characterized by increased motion of the β_D_ strand and surrounding loops in 1P-GAB1_N-term_-bound N-SH2^WT^. **(G)** Representative ^15^N-CEST profiles of residues located in β_C_ (Leu43) and β_D_ (Thr52) for 1P-GAB1_N-term_-bound N-SH2^T42A^ showing the lack of a second-state dip. **(H)** Few remaining residues with a broad shoulder in CEST are colored tan onto the 1P-GAB1_N-term_-bound N-SH2^T42A^ X-ray ensemble. **(I)** Comparing ^15^N-CPMG relaxation dispersion curves for the peptide-bound forms of both variants, showing full quenching of ms motion in the beta strands for the mutant form. **(J)** Only a few residues show CPMG dispersion for N-SH2^T42A^ shown in tan color. **(K)** X-ray ensemble structure of apo N-SH2^WT^ color-coded by *k_ex_* values derived from the 2-state fitting of CPMG data (see **Fig. S17** for all CPMG profiles). CPMG data fitted into two groups: *k_ex_ =* 3550 ± 320 s^-1^ (group 1, green) and 7000 ± 670 s^-1^ (group 2, magenta) with underdetermined populations. Tan-colored residues don’t fit into a group. **(L)** Apo N-SH2^T42A^ CPMG was fitted into two groups: *k_ex_ =* 1260 ± 330 s^-1^ (group 1, green) and underdetermined *k_ex_* > 10000 s^-1^ (group 2, magenta) with underdetermined populations (see also **Fig. S18** for CPMG profiles), while residues that don’t fit to a group are colored tan. In all structure figures, black denotes prolines or residues overlapped in ^1^H-^15^N HSQC spectra.

Analysis of CPMG profiles (R_ex_ > 1 s^-1^) for the same sample using first a two-state exchange model identified three distinct groups of residues, based on exchange rates (*k_ex_*) (**Fig. 5C, 5E, Fig. S14**). Group 1 residues fit the slow exchange rate of 104 ± 8.5 s^-1^ derived from CEST (colored in brown in **Fig. 5C, 5E**) and were predominantly in the β_C_ strand. Residues in groups 2 and 3 had faster exchange rates (**Fig. 5C**). Inspection of CPMG profiles of residues in group 2 (**blue in Fig. 5E**) revealed six probes that required fitting to a three-state process (B↔A↔C, **Fig. 5D**, **S15**), the slow process detected in CEST plus a faster process with a *k_ex,AC_*of 2600 ± 100 s^-1^ (**Fig. 5F scheme**). These residues were located on the outer strand β_D_, the strand that zips and unzips, as well as in EF and DE loops. In addition, the CPMG data of Ile54, which exhibited a slower exchange rate (*k_ex,AB_*) of 45 ± 5 s^-1^ in CEST analysis, also needed to be fitted to a three-state exchange model with an additional fast A↔C interconversion rate (**Fig. S15B**). Note that His53 and His84, group 3 residues, are fitted to a 2-state fast exchange (magenta, Fig. 5C, E, S14C), which we interpret as a local fast process around these two histidines that is not related to the key process of zipping/unzipping. Although two-state fits gave visually reasonable curves, they could not account for the small but systematic deviations at low CPMG frequencies caused by the slow A↔B process independently established by CEST. Because CPMG profiles are dominated by the faster exchange contribution, a two-state model cannot simultaneously describe both processes. Therefore, we used a three-state scheme in which the slow A↔B parameters were fixed to CEST-derived values, allowing reliable extraction of the fast A↔C rate while acknowledging that the fast-exchange populations remain underdetermined. This approach provides the minimal model needed to describe the data without over-fitting.

Strikingly, in 1P-GAB1_N-term_-bound N-SH2^T42A^, neither the slow nor the fast exchange processes observed in WT are detected by CEST or CPMG (**Fig. 5G-J**). The only residues in 1P-GAB1_N-term_-bound N-SH2^T42A^ with CPMG dispersion are located on the BG loop and are undergoing a fast exchange process with *k_ex_* >3000 s^-1^ (**Fig. S16**). In light of this major difference seen in the CEST experiments of activator bound N-SH2^T42A^ versus N-SH2^WT^, combined with the major structural difference seen in the corresponding X-ray ensembles, we hypothesize that the shift of equilibrium to the fully zipped state in the mutant to be likely the major contributor to the measured increased activator affinity (see model below).

In summary, two concerted exchange processes (**Fig. 5F scheme**) were identified for the 1P-GAB1_N-term_-bound N-SH2^WT^. The first is a slow process (A↔B) defined by the transition between a major conformation A (unzipped state) and a minor conformation B (the zipped state). The second is a much faster process (A↔C) involving the transition from the major conformation A (unzipped state) to a different minor conformation C (unzipped* state) that is much faster, characterized by increased mobility of the β_D_ strand and the surrounding loops. The population of C is underdetermined because A↔C is fast on the NMR-time scale.

In an effort to “structurally” connect the slow CEST exchange process to the β-strand zipping/unzipping observed in the X-ray ensemble refinement, we examined the chemical shifts of the minor and major states from CEST. Many residues near Thr42 experience substantial local chemical-shift perturbations from the mutation itself and therefore cannot be used, however four residues provide usable probes: D40, F41, I56, and Q57 which are the key residues for zipping/unzipping (Fig. 3F-K). For these residues, the CEST minor state ^15^N chemical shifts in WT-bound closely match the directly observed ^15^N chemical shifts of T42A-bound, which is fully zipped in the crystal structure (F41: 116.5 vs 116.4 ppm, I56: 122.1 vs 122.4 ppm, D40: 119.0 vs 118.8 ppm, Q57: 128.8 vs 128.7 ppm). This correspondence supports the interpretation that the CEST minor state in WT represents the zipped conformation, while the major state corresponds to the unzipped form seen in the X-ray ensemble. Because the structural differences between these states are subtle (sub-Å backbone displacements), and many positions near T42 experience mutation-induced perturbations, these four residues represent the clearest available probes. Thus, while chemical shifts are not accurate enough to calculate those small structural differences, the CEST-derived shift comparison to T42A are consistent with the zipped/unzipped model inferred from crystallography. We therefore interpret the zipping transition as a model supported by both NMR and crystallographic data.

For the apo proteins, CEST did not detect a slow exchange, and only fast local exchange was observed by CPMG within sensitivity limits. CPMG analyses reveal similar exchange processes in both forms. Overall, the apo proteins are highly dynamic with several processes (**Fig. 5K-L, Fig. S17-S18**) being detected. Residues that were group-fitted to a 2-state exchange model are colored in green and magenta only undergo fast exchange. We hypothesize that the group 1 residues (green in Fig. 5K,L) possibly report on the Tyr66 side chain reorientation seen in the X-ray structures (Fig. S19). Additional residues showed complex dispersion that could not be group fitted.

### Proposed multi-step peptide binding mechanism

Based on the combination of NMR relaxation data and X-ray crystallographic ensembles, we propose a multi-step binding mechanism summarized in **Fig. 6**. While our structural and NMR data define the conformational states sampled during peptide engagement and timescales, they do not resolve the kinetic order of transitions among these states. We therefore present a model consistent with the observed populations and structural ensembles, without assigning a unique mechanistic pathway. In the apo state, the N-SH2 predominantly populates the unzipped β-sheet state with Tyr66 blocking full peptide engagement by occupying the peptide binding cleft. Initial phosphotyrosine binding appears to favor the unzipped conformation, based on crystallographic evidence. Our crystallographic ensemble for the p-Tyr-bound state reveals an unzipped β-sheet conformation, indicating that p-Tyr engagement can occur prior to β-sheet closure. We therefore include this p-Tyr-bound unzipped state in the mechanistic scheme because it is experimentally observed by crystallography. We thus treat it as a plausible structural intermediate rather than a required kinetic step.

**Fig. 6.**
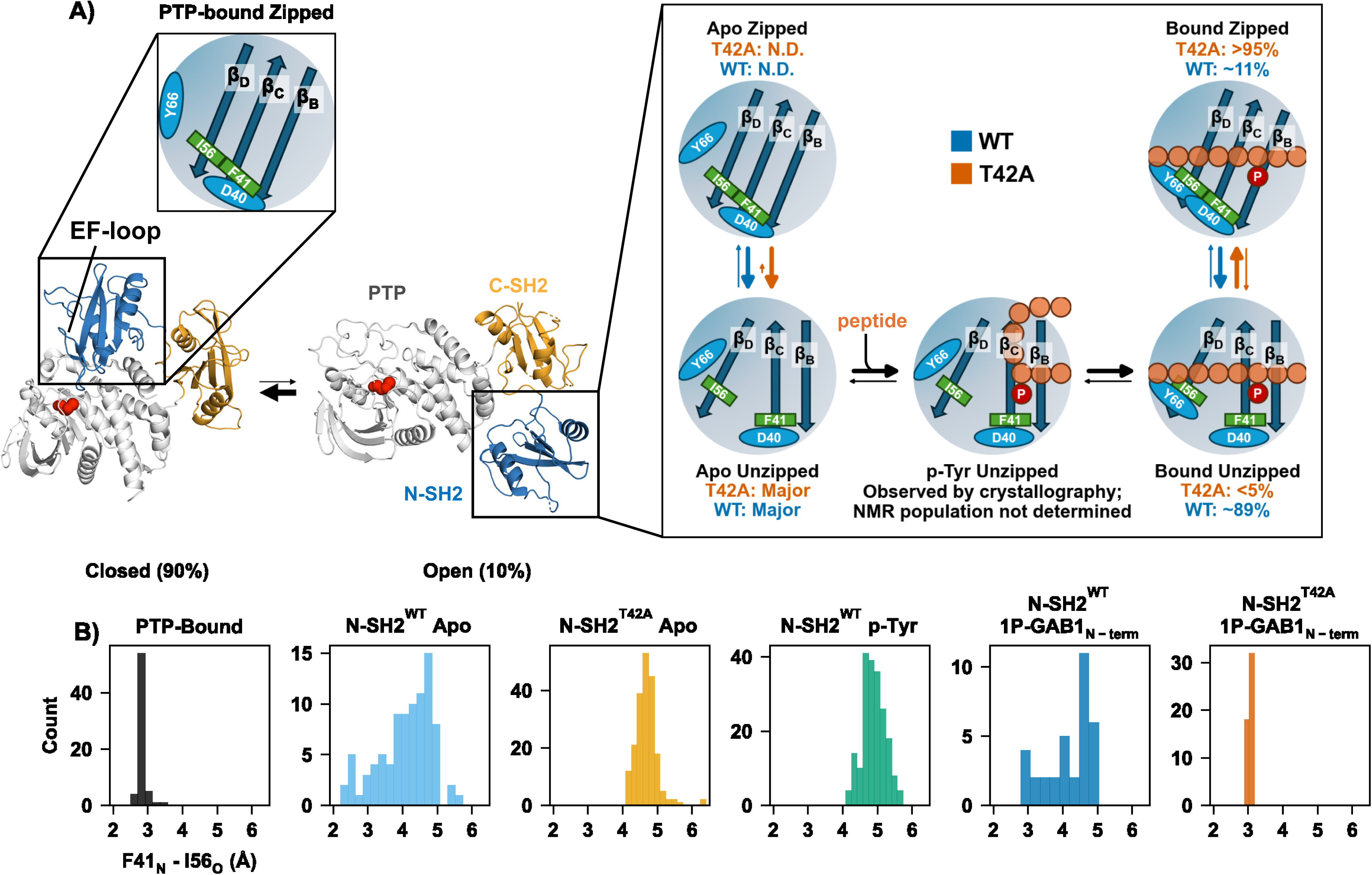
Proposed mechanism of SHP2 activation based on X-ray structures and NMR dynamics. **(A)** The N-SH2 domain, when bound to PTP, has the beta strands in the zipped conformation, Tyr66 does not block the peptide-binding cleft as in the N-SH2 apo conformation, but the EF loop causes a steric clash for full binding of phosphopeptide. When dissociated from the PTP domain, the N-SH2 is predominantly in the unzipped state as determined by X-ray ensemble, NMR dynamics within the detection limit could not detect this equilibrium. Tyr66 blocks the hydrophobic peptide-binding cleft in the apo state. In contrast, the p-Tyr binding pocket is observed to be open for p-Tyr binding. X-ray structure of N-SH2^WT^ bound to p-Tyr shows unzipped conformation. Rearrangement of Tyr66 relieves blocking and allows for binding of the peptide residues N-terminal to p-Tyr (bound unzipped). Importantly, the final step is an induced fit step from the unzipped to the zipped conformation, as illustrated by the distance between Ile56_O_ - Phe41_N_ (β_D_ to β_C_ strand) for each X-ray ensemble structure in **(B)**. This induced fit equilibrium is shifted to the unzipped conformation for of N-SH2^WT^, while N-SH2^T42A^ fully stabilizes the zipped form, characterized by hydrogen bonding between Phe41_N_ and Ile56_O_, and is further stabilized by a short hydrogen bond between the side chains of Asp40 and Tyr66. The exchange rate for this key step of zipping and unzipping is about 100 s^-1^, and the shift in populations explains the observed overall increase in peptide affinity to N-SH2^T42A^. Experimentally determined populations for the slow A↔B process in the peptide-bound WT state are ∼89% for the unzipped state and ∼11% for the zipped state from CEST analysis. In T42A, no minor state is detected in CEST (p < 5%), consistent with stabilization of the zipped β-sheet ensemble.

Tyr66 repositioning is needed to vacate the binding cleft, allowing the peptide’s N-terminal residues to fully bind, resulting in the fully bound still unzipped conformation. Importantly, we observe a subsequent induced fit step of zipping of the β-sheet, wherein a hydrogen bond between Phe41_N_ and Ile56_O_ stabilizes the extended strand conformation. The zipped state also enables the formation of a short hydrogen bond between Tyr66 and Asp40. This induced fit step is directly observed in our NMR relaxation data as the slow process (*k_ex,AB_* of ∼100 s^-1^). Strikingly, this equilibrium is shifted to the unzipped state for 1P-GAB1_N-term_-bound N-SH2^WT^, but far shifted towards the zipped state for the cancer-causing T42A mutation. This shift toward the zipped state provides a consistent explanation for the increased affinity for the activators for N-SH2^T42A^ (Fig. 2B). When the two strands are unzipped, particularly β_D_ exhibit increased mobility due to the loss of hydrogen bond between Phe41_N_ and Ile56_O_, enabling an additional faster exchange process (*k_ex,AC_*of about 2600 s^-1^).

## Discussion

SHP2 activity is tightly regulated by its N-SH2 domain, which acts as an allosteric switch between active and inactive states.(4) By keeping SHP2 inactive in the absence of appropriate signals, the N-SH2 domain plays a critical role in maintaining cellular homeostasis. Structural studies have captured the N-SH2 in both its inactive (PTP-bound)(4, 5, 41) and active (phosphopeptide-bound) conformations (31, 42), but these only represent the end states. Our study fills this gap, revealing how phosphopeptide binding drives SHP2 activation through a multi-step conformational transition (**Fig. 6**).

Given the N-SH2’s central role as SHP2’s allosteric switch, multiple studies have explored its activation mechanism.(4, 5, 32–34, 43) Our results challenge these models and, for the first time, explain how the disease-causing T42A mutation drives SHP2 hyperactivation. A previous crystal structure of the apo N-SH2 (1AYD)(44) has served as the reference model for multiple molecular dynamics studies.(32, 33) In our initial analysis, we interpreted 1AYD as resembling the peptide-bound conformation due to the presence of an unmodelled additional electron density in the peptide-binding site and the Tyr66-Asp40 hydrogen bond. Because 1AYD sequence and crystallization condition differed from what was used here, we solved the structure of apo N-SH2 using both constructs in the same crystallization condition reported for 1AYD. We confirmed the presence of molecules interacting with the EF-loop which shifts the structure to a bound conformation when the shorter construct used in the 1AYD entry is crystallized in the presence of sulfate. In agreement with our NMR relaxation-dispersion data, these observations suggest the apo N-SH2 can sample both Tyr66 conformations, and small differences in the construct or crystallization conditions may influence which state is captured (**Fig. S19**).

Our X-ray ensemble models combined with NMR dynamics experiments deliver a coherent multi-step binding mechanism for SHP2 activation (**Fig. 6**). This mechanism resolves the puzzling increase in activity for the T42A mutation, in contrast to published proposals.(32) As expected, T42A removes the Thr42/p-Tyr hydrogen bond, but the smaller alanine side chain relieves steric hindrance, increasing H-bond strength of other protein residues to the incoming phosphate, thereby slightly enhancing p-Tyr binding. A major contributing factor to the increased affinity in T42A is stabilization of the zipped β-sheet conformation in the peptide-bound state, as supported by both the crystallography and NMR. This arises from stabilizing interactions within the SH2 domain: the formation of hydrogen bonds between the backbone of Phe41 and Ile56 and between the side chains of Tyr66 and Asp40, which shift the equilibrium toward the zipped conformation. Our dynamics experiments reveal that the zipping/unzipping transitions, the key regulator for this high-affinity state, occur on the slow millisecond timescale. By vastly favoring the zipped conformation, T42A enhances phosphopeptide affinity without altering basal activity, providing a molecular explanation for SHP2 hyperactivation.

Our results challenge prior MD-based models, particularly those by Anselmi and Hub.(32, 45) Their simulations suggested that apo N-SH2 favors the zipped (termed β) conformation, while phosphopeptide binding stabilizes the unzipped (termed α) state. In contrast, our experimental data show the opposite: the apo N-SH2 is in the unzipped state, and phosphopeptides stabilize the zipped state, particularly in T42A. Anselmi and Hub’s free energy calculations incorrectly predicted reduced binding affinity for T42A, directly contradicting their conclusion that the α state (unzipped) has higher peptide affinity than the β state (zipped). Considering our measured slow millisecond exchange between zipped and unzipped states, the limited timescale of their 1 μs MD simulations could not capture this transition.

Similarly, Marasco et al.’s MD simulations combined with NMR residual dipolar coupling (RDC) experiments proposed the same model of a closed β-sheet (zipped) apo conformation(46) as in (32, 45). In fact, we would interpret their reported RDC data differently: experimental RDCs for Gly39, a probe for the β-sheet zipping, in the apo state deviate from predicted values for a fully zipped conformation as interpreted in (46), but align well when the unzipped conformations we discovered here are incorporated into the ensemble.

Using MD, Vilmmeren et al. proposed that the T42A mutation enhances phosphopeptide binding by repositioning p-Tyr deeper into the binding pocket and altering Lys55 interactions with the ligand.(35) However, we find no significant Lys55 interactions in peptide binding in our structures. Our model, in addition to the stabilizing interactions already described, shows that this enhanced stabilization occurs through an induced fit mechanism that strengthens interactions upon binding of the entire peptide, rather than solely at the phosphotyrosine-binding site. Lastly, Vilmmeren et al. proposed that the T42A mutation rewires ligand specificity.(35) We interpret their insightful quantitative published data differently: T42A enhances the N-SH2 domain’s affinity for peptides with features already preferred by SHP2, such as acidic residues at +2 and small hydrophobics at −2, which are characteristic of its native binding partners.(11) Native ligands exhibited the largest affinity increase, suggesting that T42A lowers the activation threshold for SHP2’s natural pathways rather than expanding its interaction network to non-native ligands. By stabilizing this induced fit step, T42A heightens sensitivity to native signals, prolonging SHP2 activation.

Our data further highlights the role of phosphorylation in phosphotyrosine signaling networks. The phosphate group acts as a key recognition element, increasing the affinity for the GAB1_N-term_ peptide 18,000-fold. Similarly, p-Tyr alone binds 1000- to 4000-fold weaker than the full phosphorylated peptide. This enormous difference in affinities empowers an extremely selective on/off switch, ensuring signal propagation only upon tyrosine phosphorylation of the correct peptides. Numerous studies have shown similar trends across SH2 domains, with phosphorylated peptides binding in the nanomolar range, while unphosphorylated counterparts exhibit 1,000- to 10,000-fold weaker affinity or undetectable binding.(47–50) While these differences in affinity are well established, their mechanistic basis remained unclear. Our model resolves this by proposing a multi-step process: initial weak recognition at the p-Tyr pocket induces structural rearrangements that facilitate full peptide engagement, ensuring both selectivity and high affinity.

SH2 domains share a conserved binding mechanism centered on the FLVR motif and the critical βB5 arginine, which drives p-Tyr recognition. This nearly universal mechanism ensures selective binding to phosphorylated targets. Most SH2 domains employ a two-pronged strategy: p-Tyr interacts with the conserved arginine, while additional binding pockets coordinate other phosphopeptide residues. Despite this conservation, SH2 domains exhibit sequence-specific variability. While the core p-Tyr binding is conserved, variations in specificity pockets allow for binding of distinct motifs, enabling functional diversification. In the SHP2 N-SH2 domain, Tyr66 must change conformation to allow for ligand binding, whereas in the GRB2 SH2 domain, the homologous residue Trp121 remains docked in the C-terminal binding groove in the apo state and bound to its native phosphopeptide ligands. (51, 52)This structural restraint prevents ligand binding beyond the +2 position, imparting unique ligand specificity. These variations reveal how SH2 domains have evolved distinct mechanisms to regulate ligand binding.

To extend our findings to SH2 domains in general, we analyzed the β-sheet conformations of human SH2 domains in 660 entries in the SH2 database,(53) considering the distance between the residues homologous with Phe41 and Ile56 in the SHP2 N-SH2 (**Fig. S20**). This indeed revealed two dominant conformations: a zipped state (∼3 Å) and unzipped states (>3.4 Å). These conformations are observed in a variety of SH2 domains in both apo and bound forms, indicating that zipped and unzipped states are widespread in human SH2 domains and may have diverse functional implications for different proteins. However, the biological significance of these conformations and whether they affect intrinsic properties of individual SH2 domains remain open questions for future study. In N-SH2^T42A^, the natural equilibrium is altered by shifting the peptide-bound state towards a fully zipped conformation. We propose that the equilibrium between zipped and unzipped states provides an opportunity for SH2 domains to evolve altered affinity and selectivity, with, in the case of SHP2, the T42A mutation stabilizing a conformation that enhances ligand affinity.

In molecular recognition, stronger binding is not always advantageous. If higher affinity were inherently beneficial, evolution would have favored a tighter N-SH2/phosphopeptide interaction. Instead, the N-SH2 domain is finely tuned to allow efficient ligand engagement and release, enabling SHP2 to activate in response to signals followed by a reset to the inactive state. The T42A mutation disrupts this equilibrium by stabilizing the peptide bound state, lowering SHP2’s activation threshold, leading to hyperactivation. Since SHP2 regulates multiple signaling pathways, tight control of activation is crucial. This underscores a key principle: optimal affinity in signaling proteins is not the highest possible binding strength, but the affinity that allows for effective regulation. By dissecting the interplay between phosphate recognition and peptide binding, we offer a deeper understanding of how phosphorylation so effectively modulates signaling pathways and provide key insights into the mechanism of SHP2 activation.

## Acknowledgments

We thank Prof. Lewis Kay and Dr. Jeff Bonin (University of Toronto) for their assistance with setting up CPMG pulse sequences. We are also grateful to Prof. Andrew Lee (UNC Chapel Hill) for sharing recommendations for CPMG data fitting. We thank Dr. Marco Tonelli from NMRFAM for assistance with data collection. We thank Dr. Robyn Stanfield and Dr. Marc Elsliger for X-ray beamtime at NSLS II. This study made use of the National Magnetic Resonance Facility at Madison (NMRFAM), which is supported by NIH grant R24GM141526. Helium recovery equipment, computers, and infrastructure for the data archive were funded by the University of Wisconsin-Madison, NIH R24GM141526. This work was supported by the Howard Hughes Medical Institute (HHMI) to D.K. Use of the Stanford Synchrotron Radiation Lightsource, SLAC National Accelerator Laboratory, is supported by the U.S. Department of Energy, Office of Science, Office of Basic Energy Sciences under Contract No. DE-AC02-76SF00515. The SSRL Structural Molecular Biology Program is supported by the DOE Office of Biological and Environmental Research, and by the National Institutes of Health, National Institute of General Medical Sciences (P30GM133894). The contents of this publication are solely the responsibility of the authors and do not necessarily represent the official views of NIGMS or NIH.

The Berkeley Center for Structural Biology is supported in part by the Howard Hughes Medical Institute. The Advanced Light Source is a Department of Energy Office of Science User Facility under Contract No. DE-AC02-05CH11231. The Pilatus detector on 5.0.1. was funded under NIH grant S10OD021832. The ALS-ENABLE beamlines are supported in part by the National Institutes of Health, National Institute of General Medical Sciences, grant P30 GM124169.

## Author contributions

A.W.G., R.P., A.M.O., and D.K. designed research; A.M.O. and A.W.G. collected and analyzed the NMR data, R.P. and C.S. performed X-ray crystallography experiments; A.W.G., R.P., and C.S. performed activity experiments, A.W.G., C.S., R.P. and A.M.O. expressed and purified proteins, A.W.G., R.P., A.M.O., and D.K. contributed to data analysis and interpretation; and A.W.G., R.P., A.M.O. and D.K. wrote the paper.

## Competing interests

D.K. is a co-founder of Relay Therapeutics and MOMA Therapeutics. The remaining authors declare no competing interests.

